# Fluorogenic labelling for tracking extracellular vesicle nanocarriers in the brain

**DOI:** 10.64898/2026.06.17.732894

**Authors:** Arianna Rinaldi, Myriam Catalano

## Abstract

**Background:** Reliable tracking of extracellular vesicles (EVs), key biological nanocarriers in nanomedicine, remains a major technical challenge due to the limitations of conventional lipophilic dyes, including aggregation, micelle formation, and nonspecific background signals that compromise biodistribution analyses.

**Methods:** Here, we present a fluorogenic labeling strategy based on Aco-600, a water-soluble probe exhibiting a “light-on” activation in hydrophobic environments. Medium/large EVs (m/lEVs) derived from murine BV2 microglial cells were labeled and intranasally administered to adult C57BL/6 mice. EV biodistribution and brain uptake were quantitatively assessed by ex vivo fluorescence imaging on brain cryosections at multiple time points (5–1440 min), focusing on the cortex and hippocampus.

**Results:** Aco-600 labeling enabled high signal-to-noise detection with minimal background and no evidence of dye aggregation artifacts. Quantitative analysis revealed a consistent spatiotemporal distribution profile across brain regions, with peak signal intensity at 60 minutes post-administration, followed by progressive clearance. This approach provided reproducible and sensitive tracking of EV biodistribution following a clinically relevant intranasal delivery route.

**Conclusions:** Our findings establish fluorogenic labeling as a robust and artifact-minimizing strategy for in vivo EV tracking. This method enhances the accuracy of biodistribution studies and supports the development of EV-based nanomedicine platforms, particularly for central nervous system delivery applications.

## Introduction

Extracellular vesicles (EVs) is the collective term describing a heterogeneous group of lipid bilayer membrane-delimited nanoscale structures, which includes small extracellular vesicles, sEVs (<200 nm in diameter) and other medium and larger vesicles, m/l EVs (>200 nm). EVs are released by virtually all cell types into the extracellular space, and serve as key biological nanocarriers of nucleic acids, proteins, and lipids mediating intercellular communication and the transfer of bioactive molecules thereby modulating physiological and pathological processes (Théry et al., 2018). Owing to their intrinsic biocompatibility and targeting potential, EVs have attracted increasing interest as platform for nanomedicine and drug delivery, particularly for applications involving the central nervous system.

A critical barrier to the advancement of EV-based therapeutics is the lack of reliable approaches for accurately tracking of their biodistribution and cellular interactions in vivo. A wide range of labeling strategies have been developed to investigate EV fate, including genetic engineering approaches (e.g., Cre-lox and CRISPR-Cas systems) as well as imaging modalities based on fluorescence, bioluminescence, radiolabeling, immunolabeling and magnetic resonance technologies (Bao C et al. 2023). While these methods enable spatial and temporal tracking of endogenously released EVs, allowing real-time monitoring of their fate in recipient cells or in vivo systems (Boudna M et al. 2024) they are often associated with significamt limitations, such as technical complexity or interference with EV integrity and function. Each methodology presents specific advantages and limitations; therefore, the most appropriate labeling strategy should be selected according to the experimental aim.

Among these approaches, fluorescence-based labelling remains one of the most widely adopted due to its accessibility and compatibility with high-resolution imaging.

Acoerela dyes are highly water-soluble probes that display a marked increase in fluorescence upon binding to their target, driven by changes in the surrounding hydrophobic environment. Their intrinsic photophysical properties, including relatively high quantum yield (>25%) and environment-sensitive activation, make them particularly suited for accurate tracking of nanoscale biological carriers. The pronounced contrast between bound and unbound states underlies their characteristics of “light-on” mechanism enabling high signal-to-noise detection without the need for organic solvents (https://acoerela.com/).

In this study, we evaluate a fluorogenic labeling strategy based on Aco-600 for the in vivo tracking of EV biodistribution following intranasal administration. By focusing on medium/large EVs and their delivery to the brain, we aim to establish a robust and artifact-minimizing approach for spatiotemporal analysis of EV dynamics, thereby providing a valuable tool to support the development and translational application of EV-based nanomedicine platforms.

## Results

Medium/large extracellular vesicles (m/lEVs) derived from the murine microglia-like BV2 cell line were isolated as previously described (Rinaldi et al., 2024) and characterized by nanoparticle tracking analysis (NTA), confirming successful vesicle isolation with a mean diameter of 183 ± 2 nm (Fig. 1 A). Fluorogenic labelling was performed using Aco-600 (excitation: 488 nm; emission: 586/635 nm), resulting in 45.6 ± 1.9% of vesicles exhibiting fluorescence activation, consistent with probe-membrane interaction. Of note, no significant differences in particle concentration or size distribution were observed between labelled and unlabelled m/lEVs, indicating that the labelling procedure did not alter vesicle integrity or physicochemical properties (Fig. 1 B-C).

**Figure 1.**
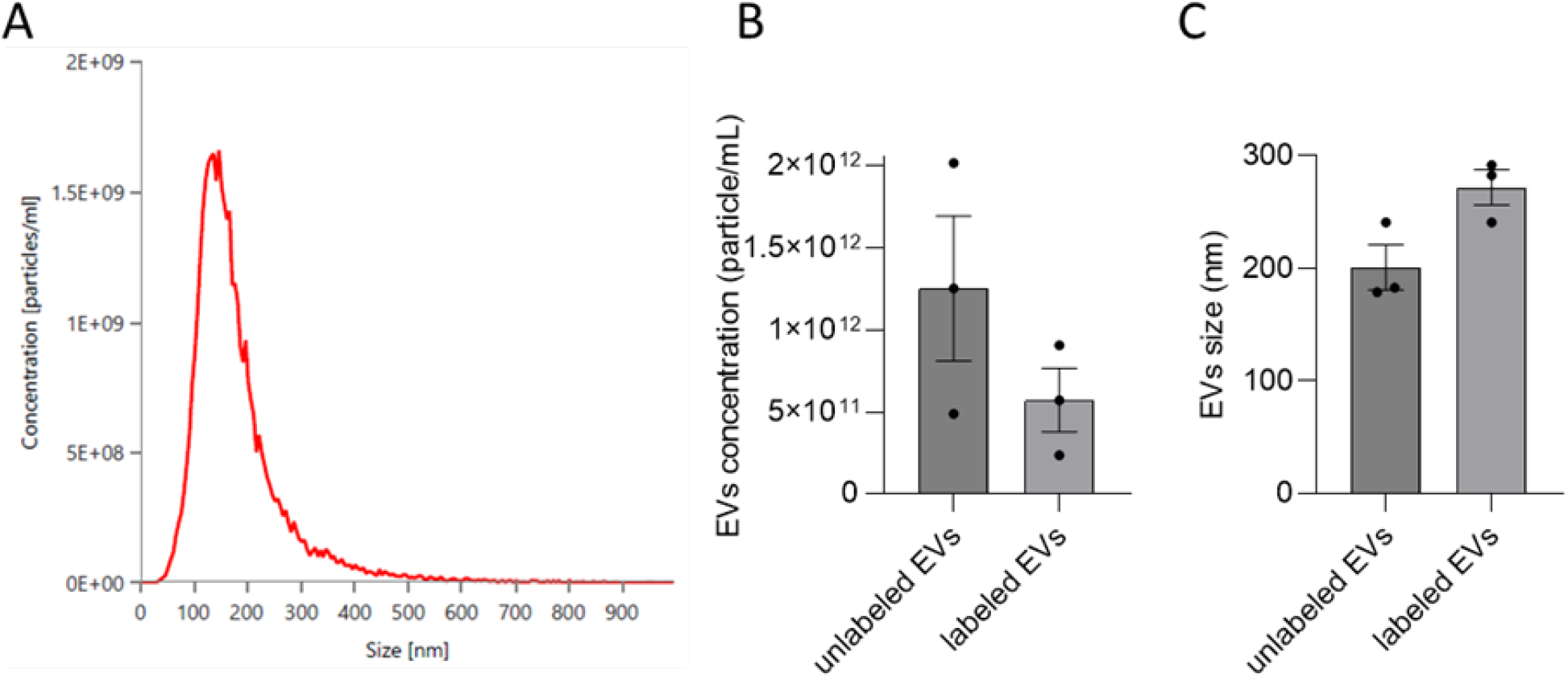
(A) Size distribution profile of m/lEVs isolated from the murine microglia-like cell line BV2, as determined by nanoparticle tracking analysis (NTA). (B) Quantification of Aco-600 negative (unlabeled) and positive (labelled) EVs. (C) Size distribution of Aco-600 negative (unlabeled) and positive (labelled) EVs.

Following intranasal administration of labelled EVs to C57BL/6 mice, vesicle biodistribution was assessed ex vivo by fluorescence imaging of brain cryosections. The presence of EV-associated signal was evaluated in the cerebral cortex and hippocampus at 5, 30, 60, and 1440 min post-administration (Fig. 2 A-B). Quantitative analysis (Fig. 2 C-D) revealed a reproducible and comparable spatiotemporal kinetic profile across both brain regions. Fluorescence intensity remained relatively stable between 5 and 30 min, followed by a significant increase reaching a peak at 60 min, and subsequently declined at later time points. While the cortex exhibited a more pronounced decrease after the peak, the hippocampus retained a relatively higher signal at later time points, without fully returning to baseline levels. This temporal profile may reflect time-dependent accumulation, interaction with target cells and subsequent clearance process within the brain tissue. The gradual decline of the fluorescence may be explained by diffusion, intracellular processing or signal degradation over time. Notably, the 60-min time point was identified as the condition showing maximal probe-target interaction, likely corresponding to optimal binding, fluorescence activation and tissue accessibility.

**Figure 2.**
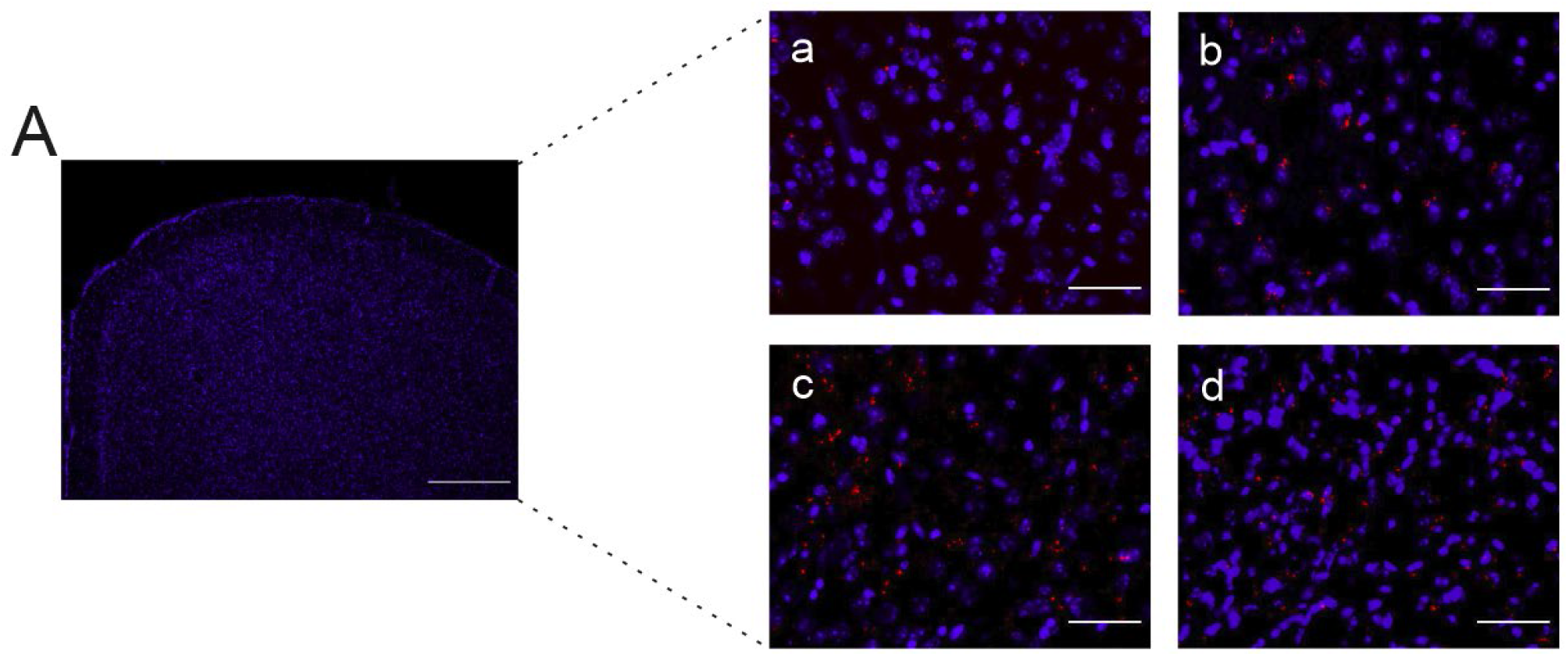
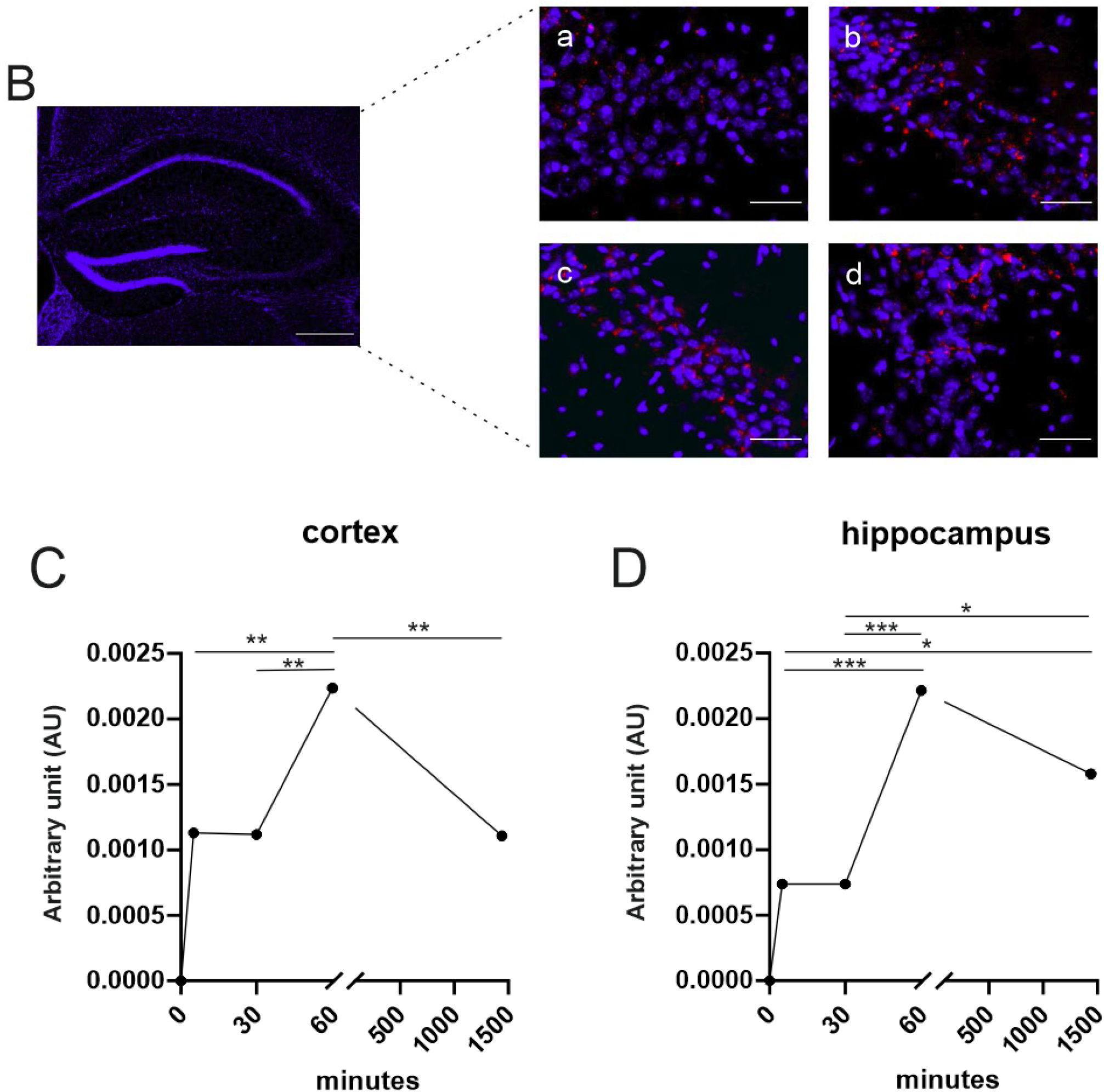
(A) Representative fluorescence images of coronal cortical brain section from mice intranasally administered with Aco-600 labelled BV2-derived EVs (red); nuclei were stained with Hoechst (blue). Scale bar: 400 μm. Right panels show higher magnification at 5 (a), 30 (b), 60 (c) and 1440 min (d); scale bar: 40 μm. (B) Representative fluorescence images of coronal hippocampal brain sections from mice intranasally administered with Aco-600 stained BV2-derived EVs (red); nuclei were stained with Hoechst (blue); scale bar: 400 μm. Right panels: magnification at 5 (a), 30 (b), 60 (c) and 1440 min (d); scale bar: 40 μm. (C-D) Quantification of Aco-600 fluorescent signal normalized to the number of nuclei in the cortex (C) and in the hippocampus (D), expressed as arbitrary unit (AU). Data are the mean ± SEM, n = 3 animals per time point. Statistical analyses were performed using one-way ANOVA followed by Tukey’s multiple comparisons test *p=0,0487, **p≤0,0071, ***p=0,0002.

## Discussion

The primary objective of this study was to establish a reliable and reproducible framework for in vivo tracking of EVs, with particular focus on brain delivery and spatiotemporal biodistribution following intranasal administration. The ability to monitor EV biodistribution in the central nervous system is is essential not only for advancing our understanding of intercellular communication, but also for supporting the development of EV-based nanomedicine and drug delivery strategies.

A growing body of evidence has demonstrated that EVs mediate bidirectional communication between brain cells and peripheral tissues, and several engineered models have been developed to visualize EV trafficking in vivo (Fordjour et al., 2022; Mathieu et al., 2021). Among these, tetraspanins such as CD81 have been extensively used as markers of EV biogenesis and transport, enabling cell-specific labelling and tracking approaches. For instance, reporter systems based on fluorescently tagged CD81 (e.g., CD81-mNeonGreen) have enabled real-time visualization of EV exchange across tissues (Fordjour et al., 2023), including long-range transfer between tumors and distant organs (Ye Y. et al 2022). While these genetic strategies provide powerful tools to study endogenous EVs, they often involve complex experimental setups and limited flexibility for exogenous EV administration studies.

In this context, a key strength of this work lies in the implementation of an experimentally accessible and in vivo-compatible pipeline that combines a minimally invasive administration route in wild-type mice with quantitative spatial and temporal analysis. Intranasal delivery represents a strategically relevant approach for brain targeting, as it enables direct access to brain regions bypassing the blood-brain barrier, avoiding gastrointestinal absorption and circumventing hepatic first-pass metabolism (Born J t al. 2002). Notably, in human subjects, intranasal administration also constitutes a painless, non-invasive and straightforward drug delivery method that is easy to manage and readily repeatable, typically associated with a rapid onset of action and a favorable tolerability profile (Arora P et al 2002). Consistent with this enhanced brain bioavailability, our data demonstrate that following intranasal administration, EV-associated signals are detectable in both the cortex and hippocampus at early time points, indicating efficient brain entry and distribution. The peak signal detected at 60 min post-administration is comparable to observations obtained with other dyes in healthy animals, where EVs display a diffuse distribution pattern and are largely cleared within 24 hours (Kodali M et al 2019).

Importantly, the identification of a reproducible kinetic profile provides a practical reference point for future studies aiming to investigate EV-tissue interactions. This temporal window likely reflects a balance between EV accumulation, interaction with resident cells, and the onset of clearance mechanisms. From a methodological perspective, defining such a peak is crucial for standardizing experimental designs, optimizing sampling time points, and improving cross-study comparability. In contrast, in murine models of disease, including stroke, neuroinflammation, tumors and neurodegeneration have been shown to exhibit prolonged EV retention and preferential accumulation in affected regions, likely driven by inflammatory and tissue-specific cues (Kodali M et al. 2019; Ma X et al 2020). In our study, the comparable behavior observed between cortex and hippocampus suggests a relatively homogeneous distribution pattern under physiological conditions, further supporting the reliability and reproducibility of the proposed approach.

A major advantage of this work lies in the use of a fluorogenic labeling strategy that minimizes nonspecific background and avoids aggregation artifacts commonly associated with lipophilic dyes. This feature is particularly relevant for quantitative in vivo studies, where signal specificity is critical for accurate interpretation of EV biodistribution and kinetics. By enabling high signal-to-noise detection without altering vesicle properties, this approach addresses a key limitation in current EV imaging methodologies. While the present study relies on ex vivo analysis, which ensures high spatial resolution and quantitative robustness, it captures EV distribution at discrete time points rather than continuous dynamics. Future integration with real-time in vivo imaging approaches could provide complementary insights into EV trafficking and kinetics. In addition, extending this framework to different EV sources, sizes and pathological conditions will be essential to assess its general applicability.

Overall, this work provides a robust and accessible methodological platform for in vivo EV tracking in the brain. By defining delivery efficiency, spatial distribution and temporal kinetics, the proposed approach supports the rational design of experiments aimed at understanding EV biology in vivo and facilitates the rational design of EV-based nanomedicine strategies for central nervous system targeting.

## Materials and methods

### Cell culture

BV2 murine microglial-like cells were maintained in Dulbecco’s Modified Eagle Medium (DMEM) supplemented with 10% heat-inactivated fetal bovine serum, penicillin (100 IU/mL), streptomycin (100 mg/mL), and amphotericin B (2.5 mg/mL). Cultures were kept at 37 °C in a humidified incubator with 5% CO_2_.

### EV isolation

Conditioned media were harvested and clarified by centrifugation at 1,000 × g for 5 min to eliminate residual cells and debris. Supernatants were subsequently centrifuged at 10,000 × g for 30 min at 4 °C to obtain medium/large EV fractions. Pellets were resuspended in 0.22 µm-filtered PBS (Thermo Fisher Scientific).

### EV fluorescent labelling

EVs were incubated by shaking with Aco-600 (Acoerela) at 2 uM for 1 h at 37° following the manufacturer’s instructions. To remove the unbound dye EVs were centrifuged at 2,500 × g for 10 min and then two-times washed in PBS (10,000 × g for 30 min). EVs were resuspend in 0.22 µm-filtered PBS (Thermo Fisher Scientific).

### Nanoparticle tracking analysis

EV size distribution and concentration were assessed using a NanoSight NS300 instrument (Malvern Panalytical) in both scatter and fluorescence modes (488-nm laser). Samples were diluted in PBS to a final volume of 1 mL to achieve 20-100 particles per frame. For each condition, five 60-s videos were recorded and analysed with NTA Software 3.4 (Dev Build 3.4.4) using an sCMOS camera, camera level 14, and detection threshold 4.

### Animals

Adult male C57BL/6J mice (2-6 months of age) were used for all experiments. Animals were group-housed (4-6 per cage) under standard laboratory conditions (temperature 20–22 °C; relative humidity 50–55%; 12-h light/dark cycle with lights on at 6:00 a.m.), with unrestricted access to food, water, and basic environmental enrichment. All procedures complied with European legislation (Directive 2010/63/EU) and were authorized by the Italian Ministry of Health (authorization no. 664/2021). Animal health and welfare were monitored throughout the study.

### Intranasal EV administration

Mice were anesthetized with isoflurane while maintaining spontaneous respiration. To enhance nasal mucosal permeability, each nostril was pre-treated with 4 µL of hyaluronidase (10 mg/mL; Sigma Aldrich). After a 30-min interval, mice received fluorescently labeled EVs. A total of 1.4 × 10^11^ EVs (6 µL per nostril), following the protocol described in Rinaldi et al. 2024, was administered intranasally.

At defined time points after EV delivery (5, 30, 60, or 1440 min), mice were perfused transcardially with ice-cold PBS. Brains were removed, post-fixed in 4% paraformaldehyde, cryoprotected in 30% sucrose, and snap-frozen. Coronal cryosections (20 µm) were rinsed in PBS, blocked for 1 h at room temperature (3% goat serum in 0.3% Triton X-100), and counterstained with Hoechst 33342 (1:1000, 1 h, RT; Thermo Fisher Scientific) to visualize nuclei. Sections were mounted using Dako Fluorescence Mounting Medium (Agilent) and imaged with a Nikon fluorescence microscope. Quantitative analyses were performed using MetaMorph 7.6.5.0.

### Statistical analysis

Data are presented as mean ± SEM. Normality was assessed using an alpha threshold of 0.05. Statistical analyses were conducted with GraphPad Prism (v9.3.1, San Diego, CA, USA). The specific tests applied to each dataset are indicated in the corresponding figure legends.

## CRediT authorship contribution statement

<u>Arianna</u> Rinaldi: Software, Methodology, Investigation, Formal analysis. Myriam Catalano: Writing – review & editing, Writing – original draft, Validation, Supervision, Resources, Project administration, Methodology, Funding acquisition, Formal analysis, Data curation, Conceptualization.

